# Juvenile Atlantic Sturgeon Survival and Movement in Proximity to an Active Cutterhead Suction Dredge

**DOI:** 10.1101/2024.03.01.582914

**Authors:** Matthew Balazik, Douglas Clarke

## Abstract

During 2019 and 2020, 268 (30-71cm fork length) juvenile Atlantic Sturgeon were captured and released in proximity to an active cutterhead suction dredge among three sites in the James River, Virginia. Of these captures, 30 were implanted with acoustic tags and telemetry was used to track their movements. Three juveniles telemetry tagged prior to this project were also detected moving within dredge operations. Cumulatively, tagged juveniles made at least 125 passes of the dredging operations with no evidence of detrimental impact in terms of survival and all were detected following the cessation of dredging. Juveniles were captured, presumed feeding, around 100m from the dredge in areas that could easily be avoided if the dredge created a stressful environment. No significant differences in catch-per-unit-effort were found when trawl catch was compared to a reference location or when monitoring gill netting catch 100m down current of a dredge over a month-long period at one of the sites. These results suggest a finding of very low risk of entrainment, migratory blockage or avoidance within 100m of an active cutterhead suction dredge by juvenile Atlantic Sturgeon.

## Introduction

Atlantic sturgeon *Acipenser oxyrinchus oxyrinchus* (ATS) is a federally protected species inhabiting the east coast of the Unites States with all six distinct population segments being listed as either threatened or endangered [1,2]. ATS are a long-lived species with a relatively high generation time making recovery from low numbers a slow process [3,4,5,6,7]. A number of obstacles to the recovery of ATS have been identified, with navigation dredging being listed as a potential threat by altering habitat and macroinvertebrate communities [5,6], hydraulic entrainment [8], and impediments to migration [9,10]. Research into how sturgeon species might be affected by dredging operations is very limited. A study by Parsley et al. in the Columbia River, Washington, suggested White Sturgeon *A. transmontanus* may be attracted to open water sediment placement areas in search of food [11]. Gill net, acoustic and trawl surveys in the St. Lawrence River suggest that juvenile ATS and Lake Sturgeon *A. fulvescens* inhabit sediment placement areas less when compared to control sites where dredge material is placed [12,13]. Reine et al. [9] used active and passive telemetry to show survival of five juvenile, 65-86 cm fork length (FL) ATS that were transported over 10km downstream and released beside an active cutterhead suction dredge (CSD) in the James River, Virginia. No evidence of injury, mortality or migratory impairment was observed. In a later study based on telemetry of 1m position accuracy, Balazik et al. [10] similarly showed that CSD had no noticeable effect on ATS migrating to spawning habitat in the James River, Virginia.

Entrainment of juvenile ATS by hydraulic trailing suction hopper dredges (TSHD) and CSD is a notable concern because fish swimming capabilities tend to increase with fish length, so smaller fish have less capability to escape hydraulic intakes [14,15]. Hydraulic riverine dredging is required to maintain legally required navigation depths of authorized federal channels. Juvenile ATS typically spend their first years within natal rivers; therefore, juvenile ATS would likely encounter dredge operations if dredging occurs in their natal river [6,7]. Although the studies cited above provide useful insights, extensive knowledge gaps persist with respect to juvenile ATS behavior to active dredging operations.

Routine maintenance dredging of the James River, Virginia, federal navigation channel occurs annually in areas that juvenile ATS (here treated as 0-2yr), are known to inhabit. To address some knowledge gaps in support of dredging project management decisions, the U.S. Army Corps of Engineers, the Engineer Research and Development Center, Virginia Commonwealth University, and Cottrell Contracting Corporation partnered to conduct an investigation of juvenile ATS behavior and survival around an active CSD working in the James River, Virginia. The goals of this study were to use telemetry, catch-per-unit-effort (CPUE) and mark-recapture data to estimate survival of juvenile ATS occupying river reaches being actively dredged and to determine if juvenile ATS show avoidance behaviors in the presence of CSD due to associated sound and sediment plumes.

## Methods

This work was carried out in compliance with guidelines set by Virginia Commonwealth University’s Institutional Animal Care and Use Committee (#AD20127) and the National Marine Fisheries Service endangered species permit (#20314-01).

### Study Area

Federal regulations state that a 91.4m wide, 7.6m deep channel must be maintained for navigation in the James River, Virginia. The study occurred at three shoaling areas of the tidal portion of the river between river kilometers 45 and 103 (Fig. 1). The three shoaling areas are referred to as Dancing Point, Windmill Point and Goose Hill. Due to chronic shoaling, CSD is required at least one if not all three locations annually. The dredging period for Windmill Point was November-December of 2019 while Dancing Point and Goose Hill were dredge September-November 2020.

**Figure 1.**
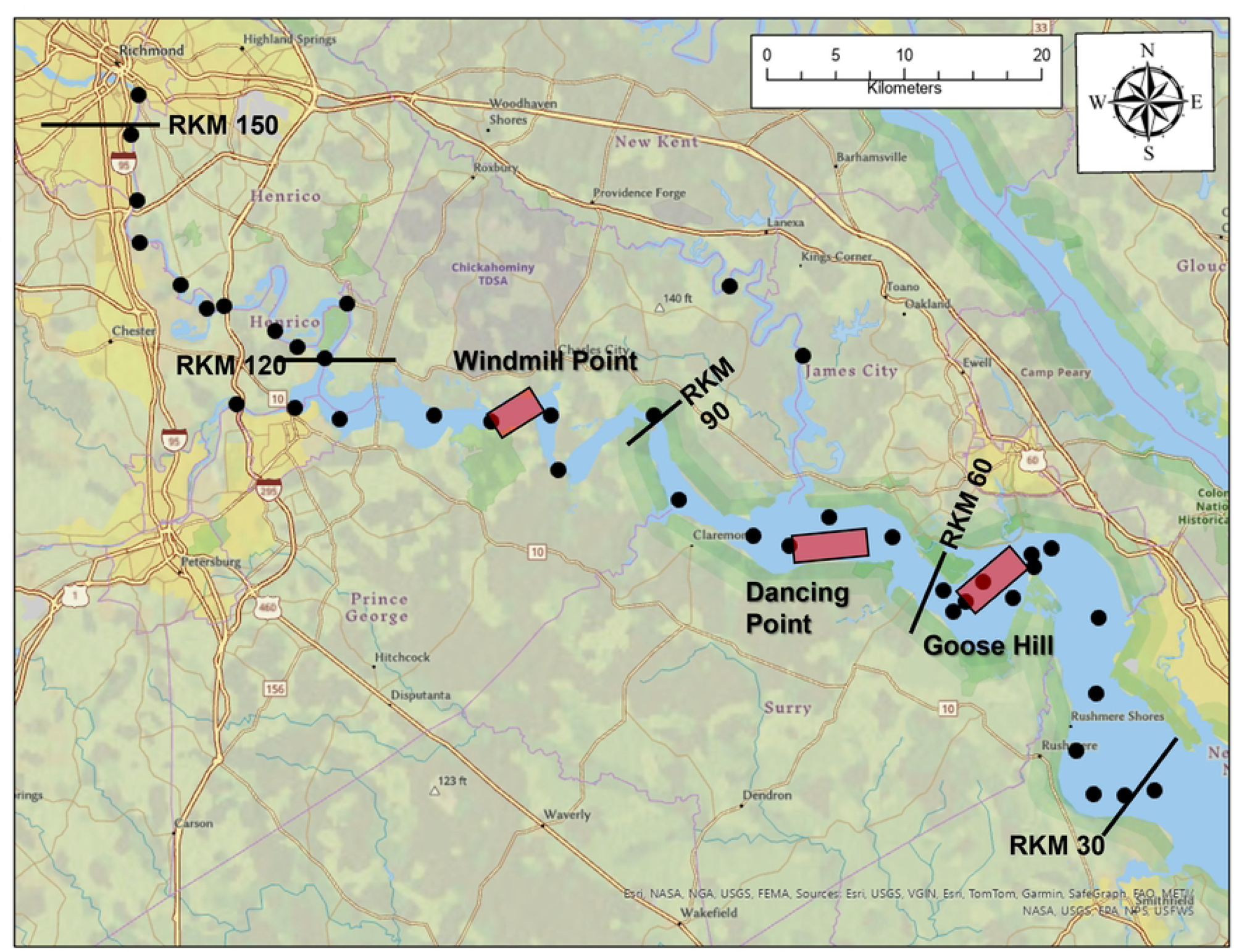
Map showing the three areas (red highlighted rectangles) juvenile ATS sampling occurred around active cutterhead dredge operations. The black points are passive receiver locations. Some of the passive receivers were damaged or lost during the multiyear tracking period.

The CSD *Marion* conducted maintenance dredging at all three study locations. The *Marion* is a 40m long, 8.0m wide, 2.0m draft barge with a 1492kW hydraulic centrifugal pump connected to a 39cm inner-diameter intake pipe located just shipward of the cutterhead. Pipeline discharge velocities were approximately 3.4m/s. All dredged material was placed back into the water via a partially floating and sinking pipeline at authorized pre-determined locations at least 400m outside of the navigation channel. Discharged sediments consisted almost entirely of silt and clays at all three locations.

### Fish Sampling

A single gill net comprising 30m mesh panels of 7.6cm, 10.2cm and 12.7cm stretch mesh was used for the study. All panels were 1.8m tall and attached in a random order for each net set. Gill nets were set perpendicular to the channel axis around 100m down current of the dredge barge (Fig. 2). Distance from the dredge was measured with a Leupold RX-800i TBR rangefinder. The cutter’s position on the bottom was estimated to be 5m in front of the dredge’s bow with some slight variation due to changes in the ladder angle during normal operation. When the net was set down current from the bow side, the gill net was about 95m from the cutterhead, whereas the net was about 145m from the cutter when set on the stern side.

**Figure 2.**
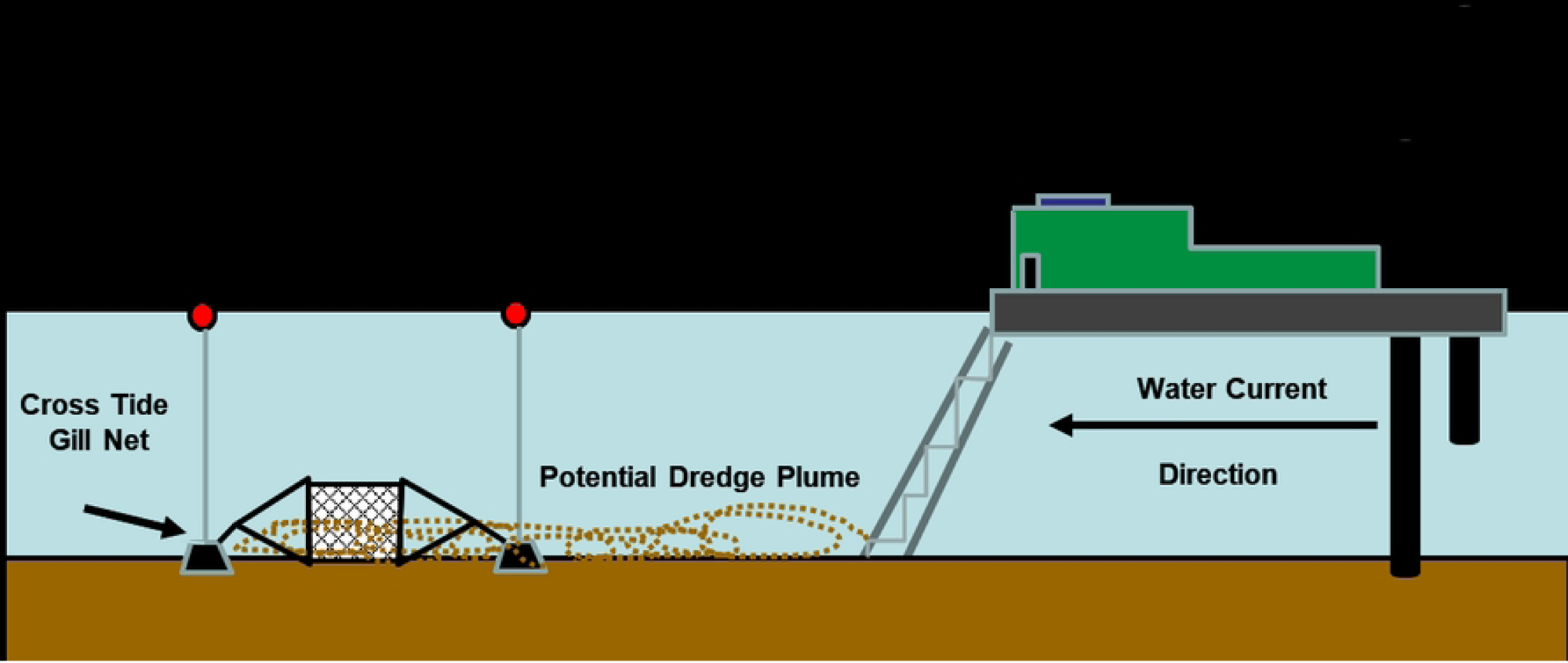
Example of how the gill nets were set in relation to the active cutterhead dredge. Not drawn to scale.

Net deployments were typically about 15min before slack current to about 15min after the water current began to flow towards the dredge after the tide changed. Net set times ranged from 40min-1hr. CPUE was total ATS caught in the net during a tide change. Nets were set in this method to see if juvenile ATS avoided the turbidity plume created by the cutterhead digging into the substrate or responded to sounds generated by the digging process. The plume would have been traveling through the sampling area for the duration of the flood or ebb tidal currents which is at least 4 hours in the James River, Virginia. All three locations dredged during this study are characterized by relatively high background turbidities and suspended sediment concentrations, which would be slightly elevated by the digging activities [16]. Discharge plumes at in-water placement areas have greater dredge-induced turbidities and suspended sediment concentrations [16], but placement areas are located well outside of the channel boundaries. Age 0-2yr ATS in the James River, Virginia, tend to confine their movements to within the comparatively deeper channel waters (Unpublished data, Matthew Balazik, Virginia Commonwealth University).

Captured ATS were scanned for a passive integrated transponder (PIT) tag. If no PIT tag was detected, a PIT and T-Tag was injected beside the dorsal fin. ATS FL was measured in cm and a fin clip was taken from the pelvic fin for genetic analysis. All fish were released at the sampling location after processing which always coincided with water currents moving toward the cutterhead.

Trawl sampling only occurred at the Windmill Point as by the time Dancing Point and Goose Hill sampling occurred, ATS had attained a size that rendered similar trawling methods ineffective. At the Windmill Point study area, a 4.9m wide SRT benthic trawl with 76cm X 38cm doors designed by Innovative Net Systems was towed with the tidal current alongside the dredge and at a reference location about 5km upstream of the dredge which is beyond the influence of the dredge plume at Windmill Point (Fig. 3). The trawl was towed by a single 30m towline connected to a bridle with 8m warp lines to the trawl doors. Tow speeds were 3.0-4.0km/h and started approximately 200m down current and stopped 200m up current after making an arc pattern around the dredge. The trawl tows were about 30-40m away when passing by the dredge. Two tows were made during each trawl sampling day, one alongside the dredge and one at the reference location. The dredge tow was completed first with the tow time being recorded. The tow time started when the 30m towline was fully taut and ended when trawl retrieval began. The same tow procedure and time was then repeated at the upstream reference location and CPUE was the number of ATS captured per tow. ATS were released where the trawl was retrieved and processed exactly as gill nets collections.

**Figure 3.**
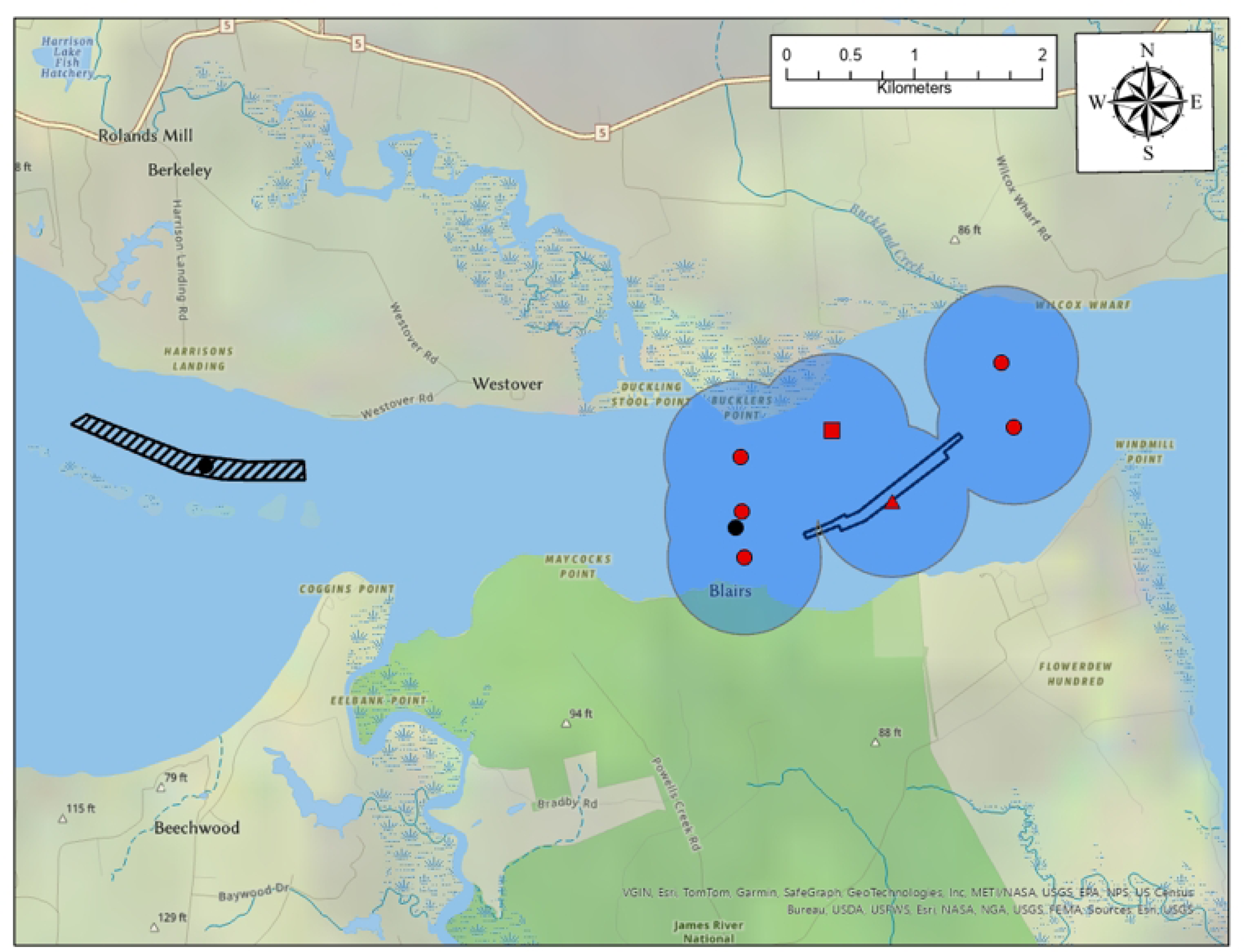
Map showing the dredge prism (black open polygon) and receiver locations at the Windmill Point study area. The black hatched polygon is the reference dredge location.

### Telemetry

ATS of suitable size were anesthetized and had a V9 Vemco telemetry tag surgically implanted using established protocols [17]. Telemetry tags with a pressure sensor providing depth information had a battery life of 214 days while the non-pressure tags lasted 912 days. All tags had a burst frequency or 20-40sec for the first 30 days then changed to 130-230sec for the remainder of the battery life. No more than three ATS were tagged during a single gill net sample. All ATS were released at the sampling location when the water current was moving towards the cutterhead. No ATS captured during trawling had telemetry tags implanted.

Telemetered ATS were tracked throughout the tidal portion of the James River using a Vemco passive receiver array (Fig. 1). Additional receivers were placed around the Windmill Point study site from November 4^th^ to December 10^th^, 2019 (Fig. 3). At Windmill Point, a receiver was attached to the stern of the dredge and another on the discharge pipe about 10m from where sediment was being placed. Range tests estimated that under prevailing weather conditions, the telemetry tags had a detection range of 600m but detections up to 1km did occur. Specific ranges tests were not conducted on the receivers attached to the dredge or discharge pipe, so sounds generated by the dredge operations may have decreased the effective detection range. However, measurements of sounds produced by CSDs working in silt/clay sediment indicate that these are relatively quiet such that an assumption can be made that detection range is not severely affected [18,19]. A receiver was attached to the dredge at Dancing Point and Goose Hill but additional receivers were not available at the time to supplement the dredge area. The receiver on the dredge was lost at Goose Hill so no detection data from the dredge are available at that location. Detection range tests were not conducted at either Dancing Point or Goose Hill, but the detection range is thought to be similar to Windmill Point.

## Results

### Windmill Point

Windmill Point dredge operations removed 129,458m^3^ of sediment from November 4^th^ to December 5^th^, 2019. The river width perpendicular to the direction of the navigation channel varied from 1.6-2.2km along the area dredged. The river cross section bathymetry of the dredged area is a confined river channel slightly wider than the federal channel surrounded by 2-3m deep mudflats. Seven gill net sets between November 4^th^ and 14^th^ captured 34 individual ATS ranging from 30cm to 47cm FL were captured by gill nets. There were no recaptures. Four sets upstream of the dredge captured 21 ATS and downstream sets captured 13 ATS (Fig. 4). These ATS are estimated to be 1yr of age. Observations of captured ATS revealed that almost all had sediment material within their oral cavity which is hypothesized to indicate feeding behavior (Fig. 5). From November 5^th^ to the 27^th^, five paired trawl tows captured 71 individual ATS ranging from 31-47cm FL (Fig. 4). The dredge trawls captured 39 ATS and the reference trawls captured 32. A two-tailed student t-test showed there was no significant difference (p=0.45) in CPUE between the reference and dredge areas.

**Figure 4.**
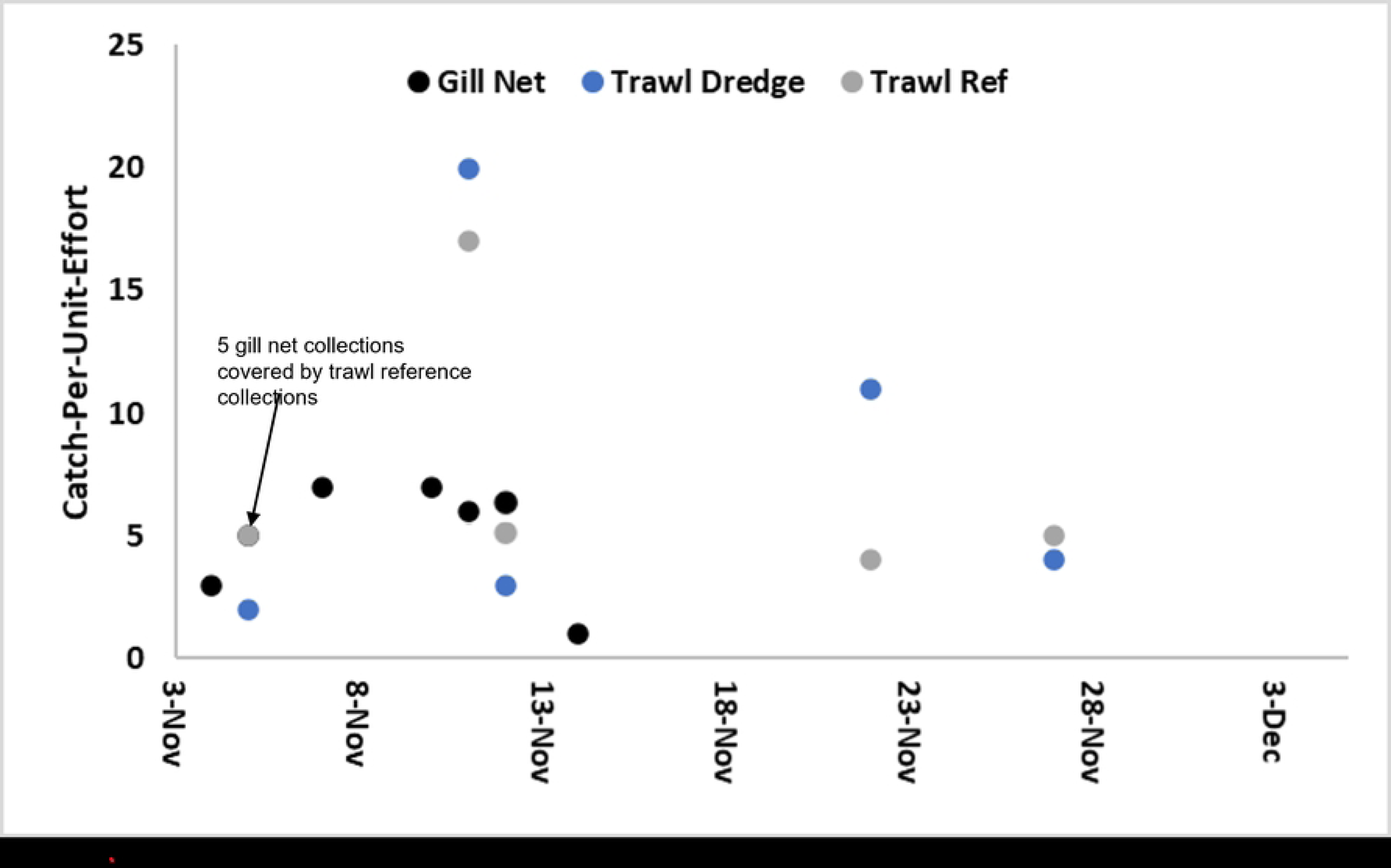
Gill net catch-per-unit-effort of ATS at Windmill Point and the trawl reference area. Note that gill nets were set on two different on November 11^th^ and that five ATS gill net collections on November 5^th^ are covered by the five reference trawl captures from the same day.

**Figure 5.**
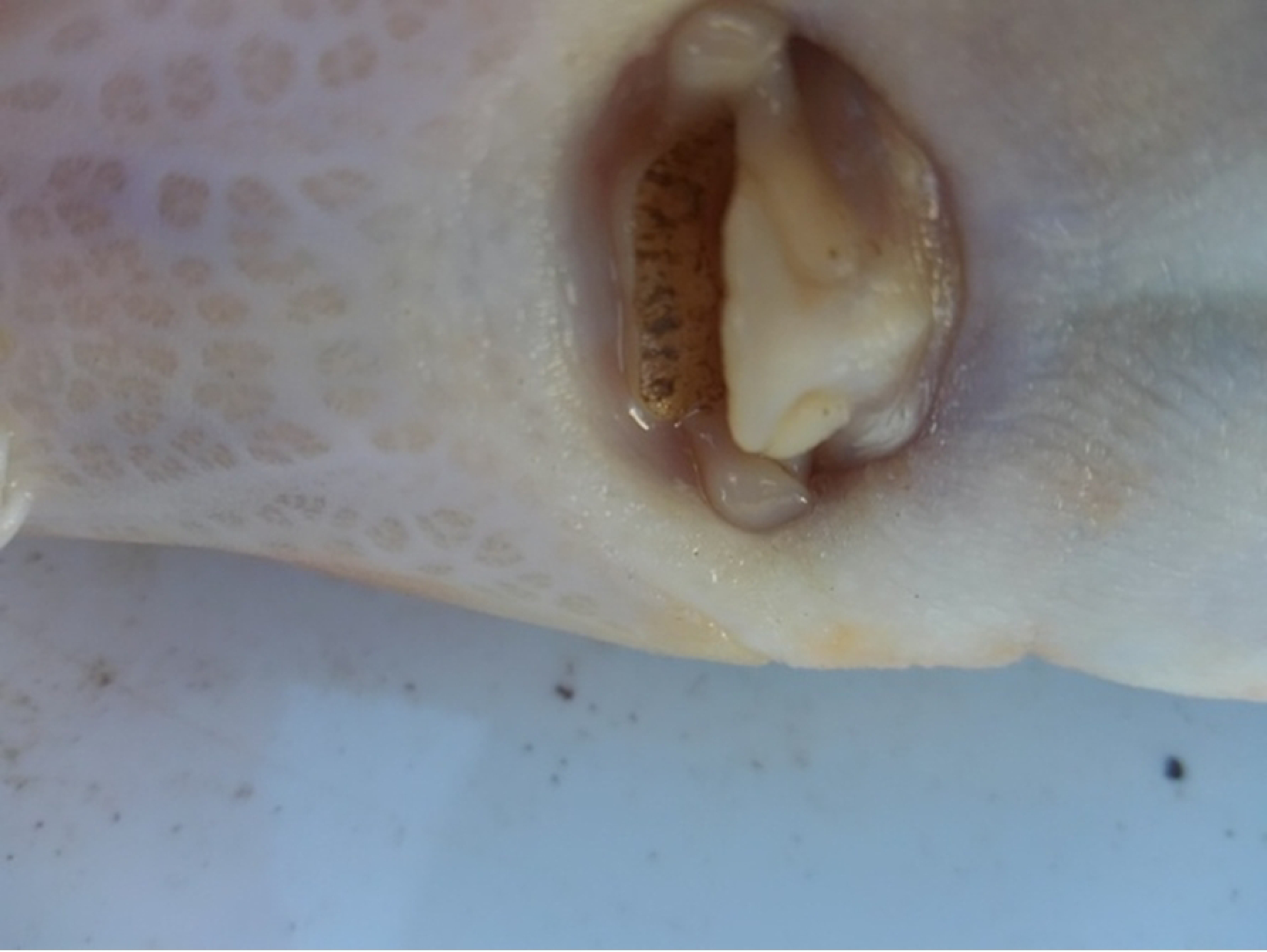
Picture showing muddy material in the mouth cavity, suggesting feeding behavior, of a juvenile ATS captured during gill net sampling.

Eighteen of the juvenile ATS captured in the gill nets were telemetered and released during Windmill Point sampling (Table 1, S1 Table). There was disparity in initial movements when telemetered ATS were released. Some immediately moved downstream or upstream after being tagged while others stayed within the dredge array (Fig. 6). Three additional ATS telemetry tagged over a month prior moved into the study area during dredge operations. The 21 tagged fish were detected within the dredge telemetry array ranging from 1-23 days totaling 199 days in the dredge array. The array recorded that tagged ATS passed the dredge a minimum of 52 times without incident with one ATS accounting for 17 passes (Table. 1, S1 Table). Data from the depth tags concurred with previous catch data suggesting that juvenile ATS spend most of time and at depths that are only available within the navigation channel (Fig. 7). The immediate post-tagging movements are likely confounded by the tagging process. Three telemetered juveniles tagged upstream more than a month prior to the dredge sampling moved through the study area and were detected by the receiver on the dredge. The best example of presumed normal behavior was VCU ATS ID 933 which was tagged 36 days prior and was within the dredge array when dredging started on November 4^th^. Fish 933 remained within the dredge array until moving downstream on November 14^th^. During the 10-day period, the fish was detected by the receiver attached to the dredge 9 days and moved passed the dredge a minimum of three times (Fig. 8).

**Figure 6.**
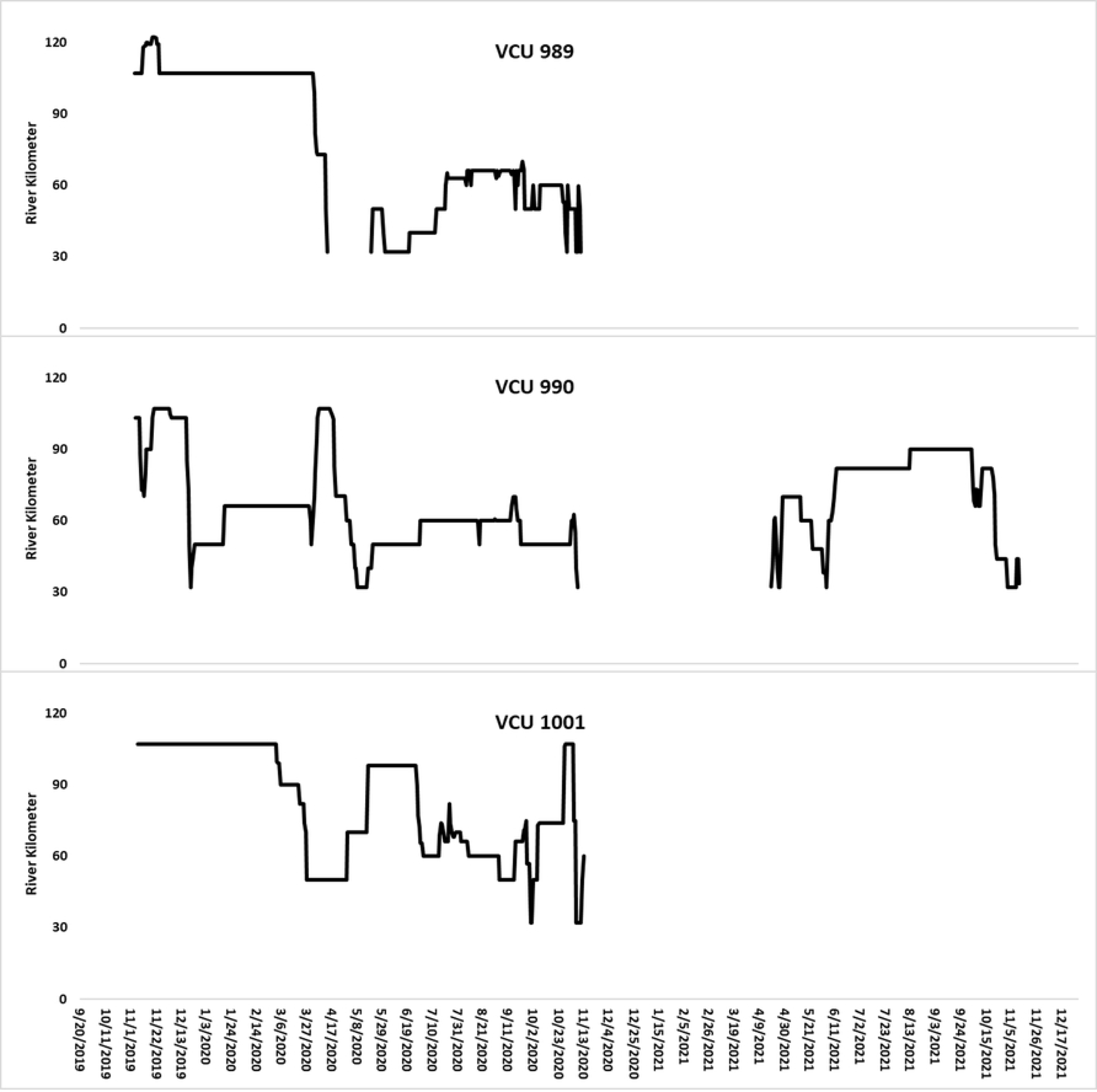
Telemetry examples of a fish moving upstream (VCU 989), downstream (VCU 990) and staying at the tagging location (VCU 1001). Blank areas are when the fish were downstream of the array within the James River.

**Figure 7.**
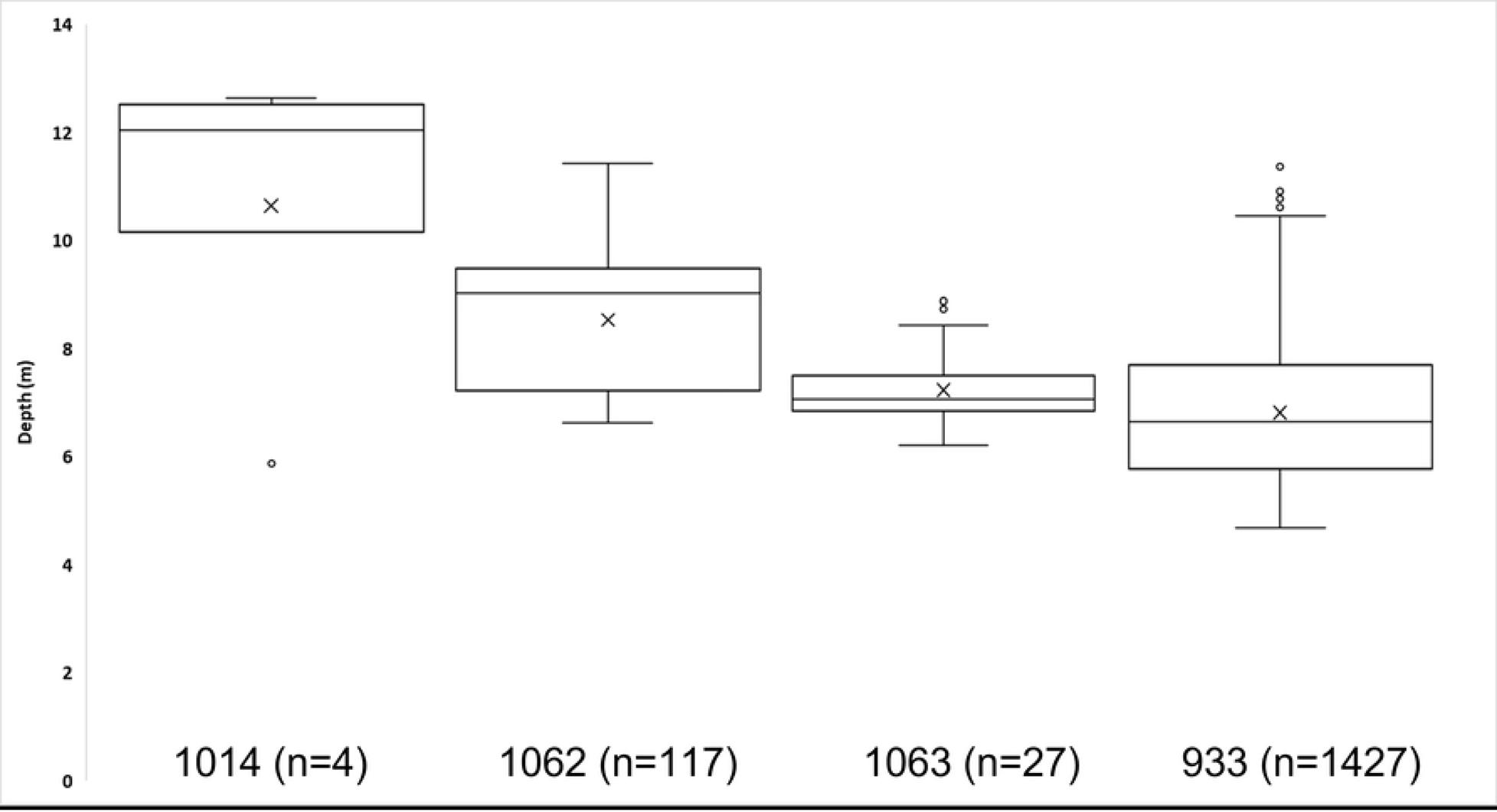
VCU ID and number of data points from the four ATS tagged with depth sensor tags.

**Figure 8.**
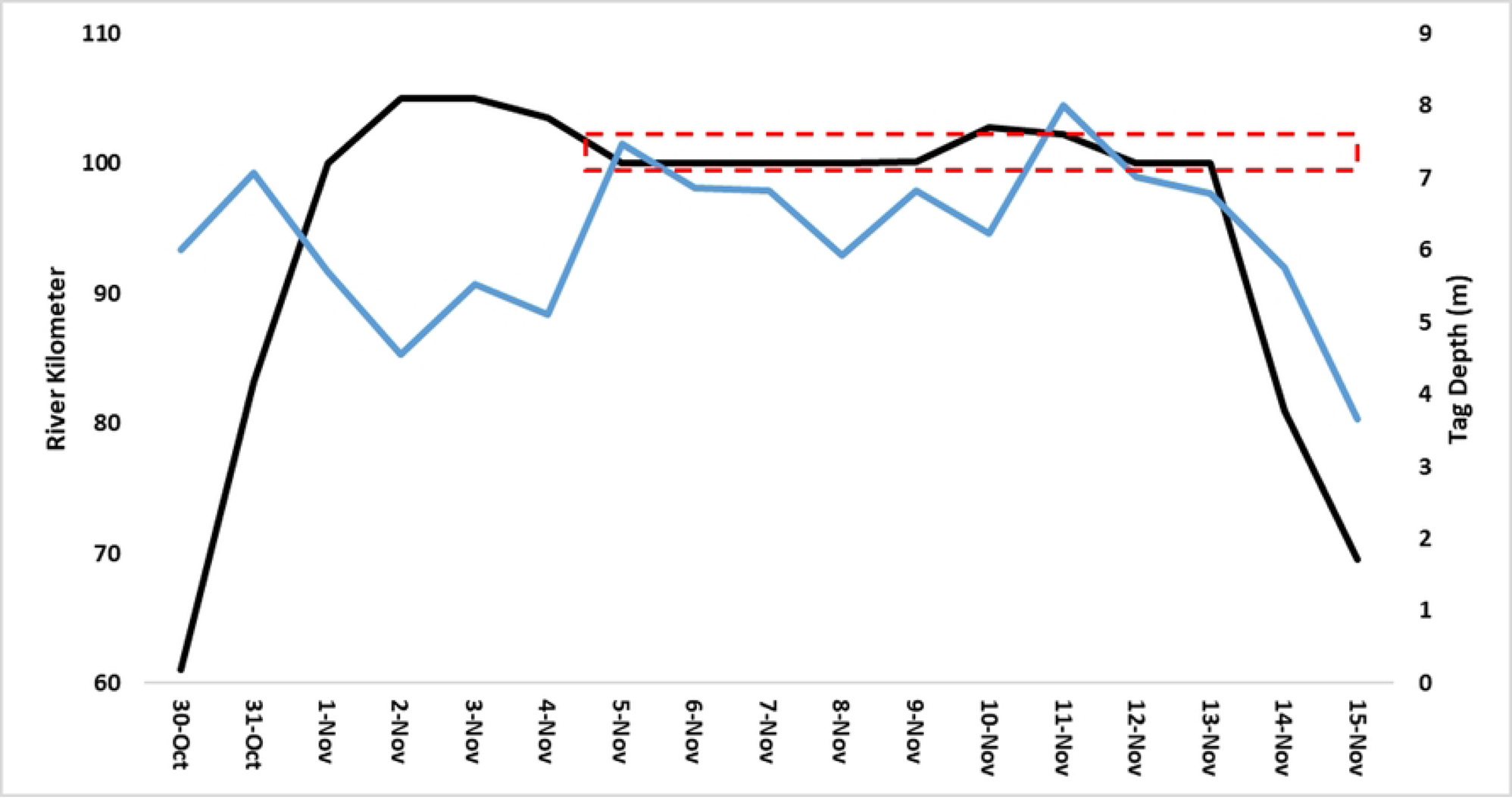
Daily average river kilometer location (black line, left axis) with daily average depth (blue line, right axis) of VCU ATS ID 933. The red dashed rectangle shows the river kilometers where dredging was occurring.

**Table 1.** Summary of telemetry data at three study locations in the James River, Virginia. The days detected in the number of days tagged ATS were detected by the receiver attached to the dredge.

| LOCATION | TAGGED JUVENILE STURGEON IN ARRAY | RELEASED | DAYS DETECTED | DREDGE PASSES |
| --- | --- | --- | --- | --- |
|  |  | UPSTREAM – DOWNSTREAM | TOTAL (RANGE) | TOTAL (RANGE) |
| WINDMILL POINT | 21 (18 NEW) | 13 - 6 | 199 (1 – 23) | 52 (0 – 17) |
| DANCING POINT | 17 (4 NEW) | 9 - 8 | 49 (1 – 16) | 31 (0 – 5) |
| GOOSE HILL | 23 (7 NEW) | 16 - 7 | NA | 43 (1 – 6) |
|  |  |  |  | 125 TOTAL PASSES |

### Dancing Point

Dredging at Dancing Point removed 220,219m^3^ of sediment from September 4^th^ to October 22^nd^, 2020. The river width perpendicular to the direction of the navigation channel varied from 4.9-5.7km along the area dredged. The river cross section bathymetry of the dredged area is a broad river channel with gradual slopes with water depths greater then 2-5m along most of the river width. Gill net sampling started October 11^th^ and ended on October 21^st^. Five upstream sets and one downstream set captured 33 and three ATS, respectively, ranging from 50-64cm FL. Four telemetry tags were deployed on the upstream side of the dredge. Information gathered from these four tagged ATL was limited as dredging was completed within four days of the ATS being released. As observed at Windmill Point, sediment was visible within the mouths of several ATS during processing.

Of the 17 telemetered juvenile ATS still inhabiting the James River, 13 (82%) were detected at the dredge and/or confirmed moving past the dredge. The previously tagged juveniles had been tagged roughly a year prior, suggesting their movements were no longer impacted by the catching and tagging processes at this point in time. The 13 telemetered ATS were confirmed moving past the dredge 28 times; however due to the distance of the passive receivers upstream and downstream of the dredge, the number of passes was likely greater. The 13 ATS were detected on a cumulative 49 days by the receiver attached to the dredge (Table 1, S2 Table). There were six instances when an ATS passed the dredge but were not detected by the receiver attached the dredge. Non-detection by the dredge receiver was likely due to the ATS passing at a distance beyond detection range as the river is over 5km wide at the dredge area. Only one ATS (VCU ATS ID 933) with a depth sensor tag was detected at Dancing Point. Depths for this ATS ranged from 1.0-6.9m deep, suggesting that the fish could have remained outside of the navigation channel even when detected by the receiver on the dredge.

### Goose Hill

Dredging and gill netting occurred at Goose Hill from October 23^rd^ to November 29^th^. Dredging removed 164,890m^3^ of sediment from the channel. The river width perpendicular to the direction of the navigation channel varied from 4.3-5.1km along the area dredged. The river cross section bathymetry resembles that of Dancing Point. Eight sets downstream of the dredge captured 48 ATS while 15 upstream sets captured 111 ATS ranging from 45-71cm FL. There were two instances when an ATS was captured twice during the Goose Hill sampling. Both were first captured on November 10^th^ and recaptured the following day. Regressions found no notable CPUE trends over the time of the sampling period (Fig. 9). Unpublished telemetry data (Matthew Balazik, Virginia Commonwealth University) indicates that most of the tagged age 2yr ATS occupied this general location in the river. Similar to Windmill Point and Dancing Point, sediment was sometimes observed within the mouth when the ATS was removed from the net.

**Figure 9.**
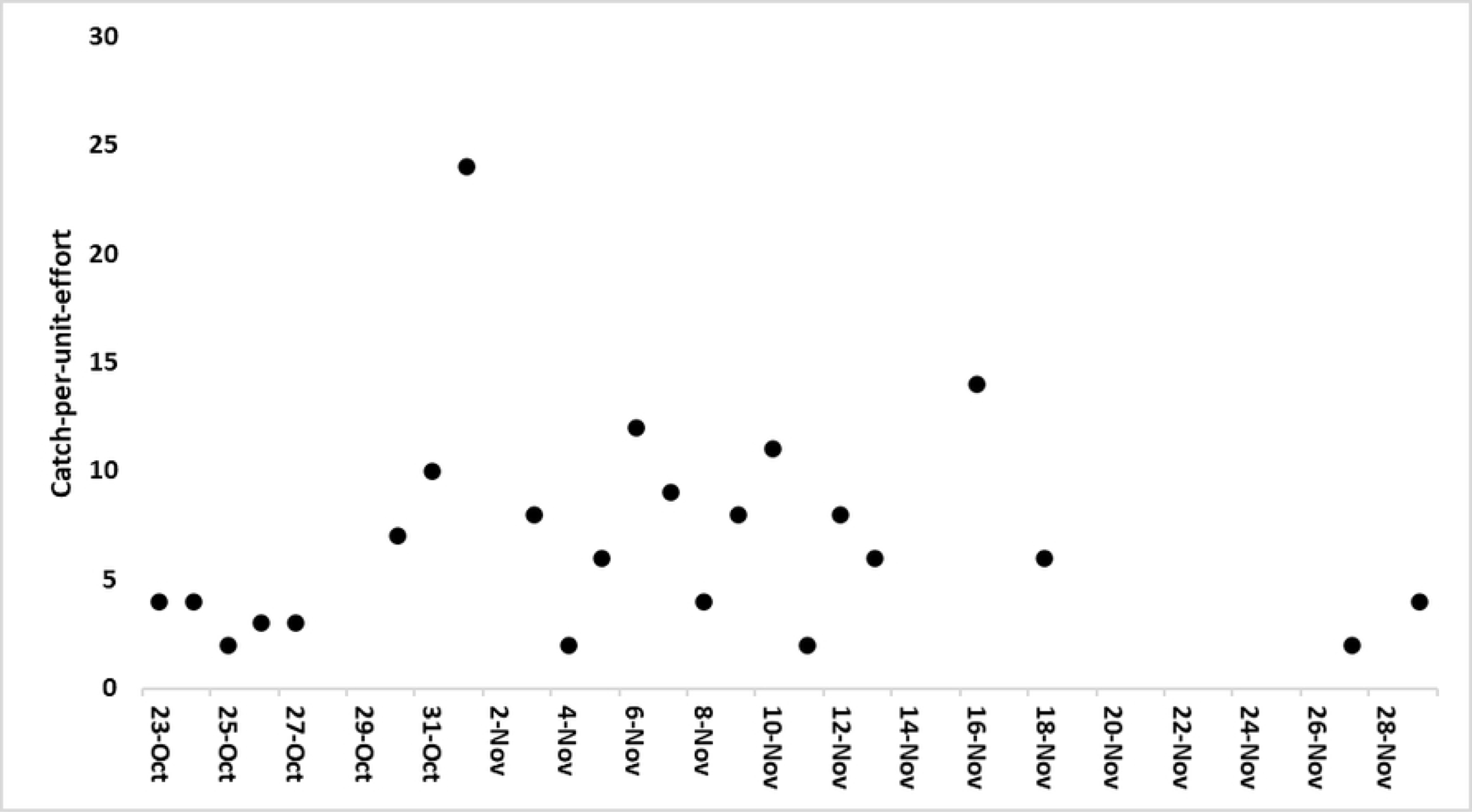
Catch-per-unit-effort at Goose Hill during the dredge period.

Seven new ATS were tagged ranging from 51-57cm FL (Table 1, S3 Table). The closest passive receivers upstream and downstream were several km away so the number of times tagged ATS moved passed the active dredge was likely greater. The seven ATS tagged at Goose Hill moved past the dredge a minimum of 13 times. Similar to Dancing Point, there were 13 (82%) ATS tagged roughly a year earlier detected in the passive array around Goose Hill. Due to the receiver attached to the dredge being lost, it is unknown how often tagged ATS were within detection range of the dredge. The ATS tagged about a year prior were confirmed to have passed the active dredge a minimum of 25 times (**Table 3**). Additionally, three of the four ATS telemetered at Dancing Point passed the active dredge a minimum of five times (**Table 3**).

## Discussion

Concerns persist that navigation dredging operations in rivers may impact migratory fishes, among them ATS. Tidal portions of rivers along the Atlantic coast are most frequently dredged by CSD. This mode of dredging entails embedding a rotating cutter in the sediment bed, an action that forms a sediment/water slurry which is hydraulically pumped through a pipeline to a designated discharge site, either upland or in-water. Descriptions of the cutterhead actions are given by Hayes et al. [20] and Henriksen et al. [21]. In the James River, Virginia, required periodic maintenance dredging is typically performed typically in this manner, with discharge back into the river outside of the navigation channel boundaries. With respect to ATS, categories of concern linked to this particular dredging process include habitat alteration, disturbance of benthic resources that represent forage items, hydraulic entrainment, and impediment or blockage of migratory movements. Concerns related to impediment or blockage of movements stem from hypothetical responses to turbidity/suspended sediment plumes created by the dredge or to sounds emitted by the dredge. The focus of the present study is on the latter topics; evidence of juvenile ATS being entrained into the cutter’s intake vortices, evidence that juvenile ATS will completely avoid an active dredge site and evidence of juvenile movements in either upstream or downstream directions.

### Hydraulic Entrainment

Concerns that CSDs could entrain and thereby injure or kill diverse aquatic organisms have proven to be difficult to confirm or refute [8]. Attention has largely focused on sea turtle and sturgeon entrainment by TSHD, which are frequently employed to maintain harbor entrance channels. Long-term systematic monitoring of take by TSHDs in the United States has documented that sturgeon can be entrained. CSD projects are not systematically monitored for take and CSD and TSHD processes are very different. CSDs are relatively stationary whereas TSHDs are mobile. A great deal of effort has been invested in systematic monitoring of sea turtle takes by TSHDs and research into dredging project management practices to reduce takes [23]. However, due to difficult logistical challenges in detecting takes by CSDs, investigators have largely relied on assessments of risk of entrainment based on knowledge of the dredging process and capabilities of the target organisms to behaviorally avoid or physically escape entrainment.

Several studies have investigated the swimming capabilities of sturgeon [23], including members of the genus *Acipenser* [24,25,26,27]. Most of these studies measured both endurance and critical burst swimming speeds in flow fields of various velocities, frequently with the objective of informing the design of fish passage structures. Other studies have specifically addressed sturgeon swimming capabilities with respect to risk of entrainment by dredges [28,29,30]. Hoover et al. [28] presented a conceptual model for determining risk of entrainment considering species specific factors including rheotaxis, sustained swimming capability, and burst swimming speed, which they treated as an escape speed. By comparing estimated CSD intake velocities of 50cm/s at 1.5m from the cutter to a measured escape speed in tandem with measures of swimming performance and behavior, they calculated susceptibility to entrainment. Juvenile Pallid Sturgeon (*Scaphirhynchus albus*) escape speed was estimated to be 51-70cm/s, which when considered with other behavioral traits led to a conclusion that they would be slightly susceptible to entrainment if they came within the influence of the intake flow field. Boysen and Hoover [29] used swimming performance to estimate risk of entrainment for trained and untrained juvenile White Sturgeon. Untrained White Sturgeon juveniles (80-82mm) had minimum escape speeds of 40cm/s, although they noted that escape speeds varied widely among individuals. Juvenile White Sturgeon “trained” by exposure to continual water flows of 10-12cm/s for extended periods had substantially higher escape speeds. Trained fish were considered to be representative of wild rather than hatchery-reared juveniles. Hoover et al. [30] used a similar approach to evaluate susceptibility to entrainment of juvenile Lake Sturgeon and Pallid Sturgeon from different populations. They concluded that the risk of entrainment for all tested cohorts was minimal unless they entered a 1.25m radius of a dredge intake flow field. Results of the above studies are consistent with results of the present study in that tracking data indicated that 1-2yr juvenile ATS moved past an active cutter hundreds of times without impingement/entrainment.

### Turbidity or suspended sediment plumes

One caveat identified by the authors of the entrainment studies described above was that in the field, additional factors could affect the outcome of dredge encounters that were not addressed in their investigations. For example, in the field sturgeon might behave differently in the presence of suspended sediments or certain dredging-induced sounds. Wilkens et al. [26] investigated survival and swimming performance of juvenile ATS exposed to suspended sediment concentrations of 100, 250, and 500mg/l for 3 days prior to swimming tests. Of 90 juveniles tested 86 (96%) survived. One fish was lost at 250mg/l and 3 at 500mg/l. Critical swimming speeds were evaluated to be moderate at 21 to 31cm/s. They concluded that the lack of substantial immediate impacts on mortality or swimming performance in the laboratory suggested that free swimming fish in the field would experience minimal ill effects from dredging operations. In regards to this field study, the juvenile ATS were not confined to the 100m area down current of dredge operations but remained, suggesting dredge operations did not generate too stressful of an environment.

### Underwater Sounds

Potential impacts of anthropogenic sound, including sounds produced by dredges, on aquatic organisms have received heightened attention in recent years. Although largely focused on the effects of intense impulse sounds, such as those associated with pile driving or underwater explosions, more subtle effects of lower intensity sound on behavior and physiology have also been identified. Relatively few characterizations of sounds produced by various modes of navigation dredging have been published. Coupled with a general lack of knowledge pertaining to the hearing capabilities of many aquatic organisms and their response thresholds to sound exposures, informed management decisions regarding the conduct of dredging operations in the presence of potentially susceptible organisms are handicapped. Sturgeon represent a prime example of the need for a better understanding of their hearing capabilities. Popper [31] submitted a report to the U.S. Army Corps of Engineers Portland District in 2005 reviewing the state of knowledge of sturgeon hearing at that time. He stated that sturgeon of the genus *Acipenser* could probably detect sounds below 100 Hz to somewhat above 1,000 Hz. Popper also suggested that sturgeon could determine the direction of sound sources but perhaps only at limited distances from the source. In a 2023 review, Popper and Calfee [32] indicated that because sturgeon lack a structural connection between the swim bladder and the inner ear, and that the swim bladder is located relatively far posterior to the ear, that sturgeon probably only detect the particle motion component of underwater sound and not the sound pressure component. Particle motion attenuates relatively rapidly with distance from the source, which implies that sturgeon may not detect sounds beyond short distances from the source. Based upon examination of limited sturgeon audiogram data, Popper and Calfee [32] reached tentative conclusions that sturgeon can detect sound from below 50 Hz to 500 to 1,000 Hz, and that sturgeon may have reduced capabilities to localize sound direction in comparison to other fishes.

Interactions between sturgeon and underwater sound can be addressed in a risk assessment framework [33,34], but technical and logistical challenges make collection of necessary input data difficult. Quantifying sound source and propagation parameters under field conditions, particularly when a dredge is involved, requires specific expertise and instrumentation. A robust risk assessment would consist of integrating hearing capabilities of the target species with characterization of the sounds of interest and knowledge of the sound exposure thresholds that would induce detrimental impacts, expressed with respect to behavioral response or injury. One limitation is that methods to measure particle motion, of particular importance for sturgeon hearing, have not been fully developed. Another limitation is that appropriate sound exposure thresholds for sturgeon have not been determined [35].

Characterization of the sounds to which sturgeon are exposed is an important part of a risk assessment. Although not conducted in the James River, underwater sounds emitted by CSDs have been characterized elsewhere [36,37,38]. Reine, Clarke, and Dickerson [36] describe the multiple sounds associated with CSD operations. Clarke, Dickerson, and Reine [37] measured sounds produced by a large CSD conducting maintenance dredging in Mississippi Sound. Reine and Dickerson measured sounds produced by a medium-sized CSD performing maintenance dredging at the Port of Stockton, California. Based on these characterizations, CSD sounds were categorized as continuous rather than impulsive, with most of the acoustic energy falling in the 70 to 1,000 Hz range. This range overlaps with the known hearing capabilities of sturgeon, so it can be assumed that juvenile ATS could hear CSD-induced sounds if they came sufficiently close to the dredge. The precise distance that juveniles would detect sounds is not predictable given available data but can reasonably be assumed to be less than 200 to 300m. Because CSD sounds are continuous rather than impulsive, sturgeon able to detect these sounds may become habituated to them relatively rapidly. A strong, persistent attraction or avoidance response would be unlikely.

Results of the present study are summarized in **Table 4** and supplement those of a previous study by Balazik et al. [10] in which survival and movements of adult ATS in the James River were examined. In that study fine spatial scale telemetry was used to follow the migration of 103 tagged adult ATS upstream to spawning habitat while a CSD was operating in the Dancing Point reach of the river. All tagged sturgeon survived and successfully reached upstream spawning habitat. Collectively 298 transits of the dredge were recorded with no indication of altered swimming speed or degree of meandering. Popper and Calfee [32] noted that Balazik et al. [10] did not collect concurrent underwater sound data, as is the case for the present study. However, in tandem the two studies provide strong evidence that the presence of an active CSD did not affect survival or movements of either juvenile or adult ATS.

## Conclusions

During the present study, 268 age 1-2yr old ATS were captured in close proximity to an active CSD. The 31 acoustically tagged ATS that were detected had 100% survival as they passed the dredge a minimum of 122 times. Juvenile ATS did not seem deterred from being within 100m of the cutterhead where the plume within the channel was likely greatest. Sediment in the oral cavities of many netted ATS suggests that ATS were feeding when caught. Juvenile ATS were still being caught towards the end of all three dredge operations even when dredging had been underway for over a month. If sound or plumes from the dredging created stressful conditions within 100m of the dredge, it is unlikely that highly mobile juveniles would be caught so close to the dredge. Although it cannot be said that dredging as conducted in the James River poses zero risk of entrainment or impeded movement, the actual risks of either impact appear to be exceedingly small.

## Acknowledgements

We would like to thank Martin Balazik, Thiwaporn Balazik, and George Trice (field assistants), Greg Garman (Virginia Commonwealth University) and Albert Spells (US Fish and Wildlife Service), Keith Lockwood for assisting and providing supplies with this project. This work was partially supported by the USACE Norfolk Division and a NOAA Section 6 grant #NA13NMF4720037, the Virginia Department of Games and Inland Fisheries award #NA13NMF4720037. This manuscript represents VCU Rice Rivers Center publication ###. The authors declare that they have no conflict of interest.

## Supporting information

S1 Table. Information for tags detected at the Windmill Point dredge location.

S2 Table. Information for tags detected at the Dancing Point dredge location.

S3 Table. Information for tags detected at the Goose Hill dredge location.

